# Spatio-temporally efficient coding assigns functions to hierarchical structures of the visual system

**DOI:** 10.1101/2021.08.13.456321

**Authors:** Duho Sihn, Sung-Phil Kim

## Abstract

Hierarchical structures constitute a wide array of brain areas, including the visual system. One of the important questions regarding visual hierarchical structures is to identify computational principles for assigning functions that represent the external world to hierarchical structures of the visual system. Given that visual hierarchical structures contain both bottom-up and top-down pathways, the derived principles should encompass these bidirectional pathways. However, existing principles such as predictive coding do not provide an effective principle for bidirectional pathways. Therefore, we propose a novel computational principle for visual hierarchical structures as spatio-temporally efficient coding underscored by the efficient use of given resources in both neural activity space and processing time. This coding principle optimises bidirectional information transmissions over hierarchical structures by simultaneously minimising temporal differences in neural responses and maximising entropy in neural representations. Simulations demonstrated that the proposed spatio-temporally efficient coding was able to assign the function of appropriate neural representations of natural visual scenes to visual hierarchical structures. Furthermore, spatio-temporally efficient coding was able to predict well-known phenomena, including deviations in neural responses to unlearned inputs and bias in preferred orientations. Our proposed spatio-temporally efficient coding may facilitate deeper mechanistic understanding of the computational processes of hierarchical brain structures.

## 1. Introduction

It is well-established that a wide array of brain areas has a hierarchical structure, including the visual system (Felleman and Van Essen, 1991; Mesulam, 1998; Harris et al., 2019; Hilgetag and Goulas, 2020). Studies have identified a link between hierarchical structures and gene expression (Burt et al., 2018; Hansen et al., 2021), suggesting that hierarchical structures are genetically determined *a priori*. Given that one of the major functions of the brain is to represent the external world (deCharms and Zador, 2000; Kriegeskorte and Diedrichsen, 2019), an ensuing question arises: How do *a priori* hierarchical brain structures attain functions to represent the external world? This question can be addressed by identifying a fundamental neural coding principle that assigns representational functions to hierarchical structures.

The traditional view of the neural coding of visual hierarchical structures is bottom-up visual information processing, whereby simple features are processed in a lower visual hierarchy and more complex features created by integrating simple features are processed in a higher visual hierarchy (Hubel and Wiesel, 1962; Hubel and Wiesel, 1968; Riesenhuber and Poggio, 1999; Riesenhuber and Poggio, 2000; Serre et al., 2007; DiCarlo et al., 2012; Yamins et al., 2014). However, this view does not consider the role of top-down pathways that are abundant even in the early visual system, such as the lateral geniculate nucleus (Murphy and Sillito, 1987; Wang et al., 2006) and primary visual cortex (Zhang et al., 2014; Muckli et al., 2015; Huh et al., 2018).

The role of top-down visual processing is especially prominent in predictive coding (Rao and Ballard, 1999; Spratling, 2017). According to predictive coding, a higher hierarchy performs top-down predictions of neural responses in a lower hierarchy. Both inference and learning of predictive coding are based on the minimisation of bottom-up prediction errors. Predictive coding has been used to explain the neural responses corresponding to prediction errors (Friston, 2005) and extends from the explanations of perception to action (Friston, 2010; Clark, 2013). A recent study combined predictive coding with sparse coding (i.e., sparse deep predictive coding) and demonstrate that it could enhance perceptual explanatory power (Boutin et al., 2021).

Nevertheless, predictive coding has several theoretical shortcomings. Since inference in predictive coding aims to minimise prediction errors, the hierarchical structure would require an additional information processing subsystem to perform this inference. In addition, because bottom-up transmitted information contains only prediction errors, predictive coding requires the presence of error units (biological neurons) in the hierarchical structure to represent this prediction error, yet such error units remain as hypothetical entities and evidence for prediction error responses is limited in some conditions (Solomon et al., 2021).

In a hierarchical structure in which information is exchanged in both directions, if the information represented by the upper and lower hierarchies at the same time is different, it is difficult to obtain a stable neural response on time domain for an external input. The reason is that if the information represented by the upper and lower hierarchy are different, different information is exchanged, and thus the information represented next time may be also different. These inter-connected structures could also produce chaotic dynamics (Rubinov et al., 2009; Tomov et al., 2014). Nevertheless, neural responses in the early visual system can be decoded as external inputs. These decodable neural responses include both neuronal spikes (Berens et al., 2012; Zavitz et al., 2016) and blood-oxygen-level-dependent responses (Kamitani and Tong, 2005; Brouwer and Heeger 2009). Therefore, it is important to find neural coding principles that enable decodable stable neural representations in hierarchical structures on time domain.

Existing efficient coding (Attneave, 1954; Barlow, 1961; Laughlin, 1981) does not require additional information processing subsystems or virtual error units, which are shortcomings of predictive coding. Unfortunately, however, existing efficient coding does not consider hierarchical structures on the time domain. Many other ideas have been proposed for representation learning (Bengio et al., 2013). Properties of the visual cortex have been successfully studied using sparse coding (Olshausen and Field, 1996). Sparse coding has been successfully implemented using artificial neural networks such as restricted Boltzmann machines for representation learning (Goodfellow et al., 2011). However, these sparse coding studies have the disadvantage that they do not take into account the passage of time, which is an important aspect in the operation of the real brain. The hierarchical structure of the brain, which ascends to the upper hierarchy from the input by the bottom-up pathway and then descends to the input by the top-down pathway again, resembles the structure of autoencoders (Bourlard and Kamp, 1988; Hinton and Zemel, 1993) of artificial neural networks. A study to obtain slow representations change using temporal coherence was implemented using autoencoders (Zou et al., 2011). However, such studies using temporal coherence lack the aspect of dynamically reacting to changes in external input or neural responses of other hierarchies.

We, therefore, propose a novel computational principle for hierarchical structures as spatio-temporally efficient coding underscored by the efficient use of given resources in both neural activity space and processing time (Fig 1). By spatio-temporally efficient coding, neural responses change smoothly but dynamically. Spatio-temporally efficient coding in hierarchical structures can also be seen as an extension of efficient coding into hierarchical structures on time domain.

**Fig 1.**
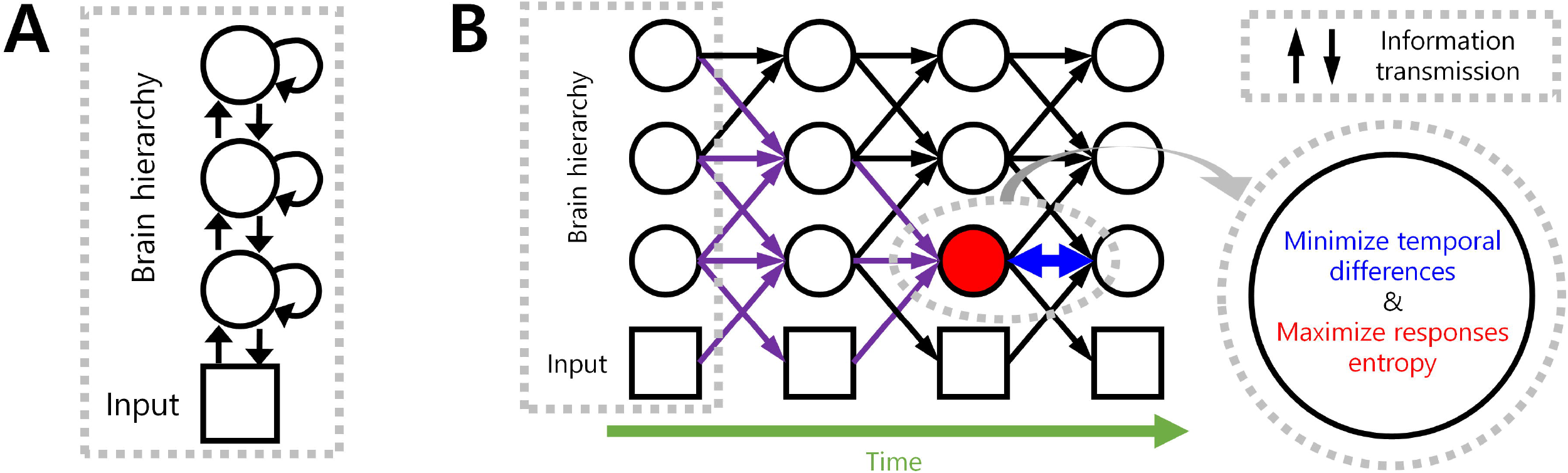
Spatio-temporally efficient coding. (A) This illustration depicts a hierarchical structure of the brain. Open black circles indicate an ensemble of neuronal units in each hierarchy of the brain. Open black square indicates visual input. Back arrows indicate information transmissions of bottom-up (upward arrow), recurrent (loop arrow), and top-down (downward arrow). (B) This illustration depicts a hierarchical structure of the brain and learning objectives. The hierarchical structure learns to represent input from the external world, which is depicted as black squares (e.g., visual input). Open black circles indicate an ensemble of neuronal units in each hierarchy of the brain. Inference based on spatio-temporally efficient coding is made by neuronal units as bottom-up, recurrent, and top-down information transmissions over time (black arrows). Learning in spatio-temporally efficient coding consists of two objectives: minimising the temporal differences between present and future neural responses and maximising the informational entropy of neural responses. For example, information transmissions (purple arrows) are optimised to minimise the temporal differences between present neural responses at the corresponding hierarchy (red filled circle) and future neural responses (circle to the right of the red filed circle) while concurrently maximising the informational entropy of neural responses at the corresponding hierarchy (red filled circle).

Spatio-temporally efficient coding enables rapid stabilization of neural responses and smooth neural representations. Through simulations, these properties (smooth temporal trajectory of neural responses, smooth neural representations, decodable stable neural responses; Fig 2, 3, and 4) and predictable phenomena (deviant neural response for unlearned inputs, orientation preference bias; Fig 6, 7 and 8) were confirmed.

**Fig 2.**
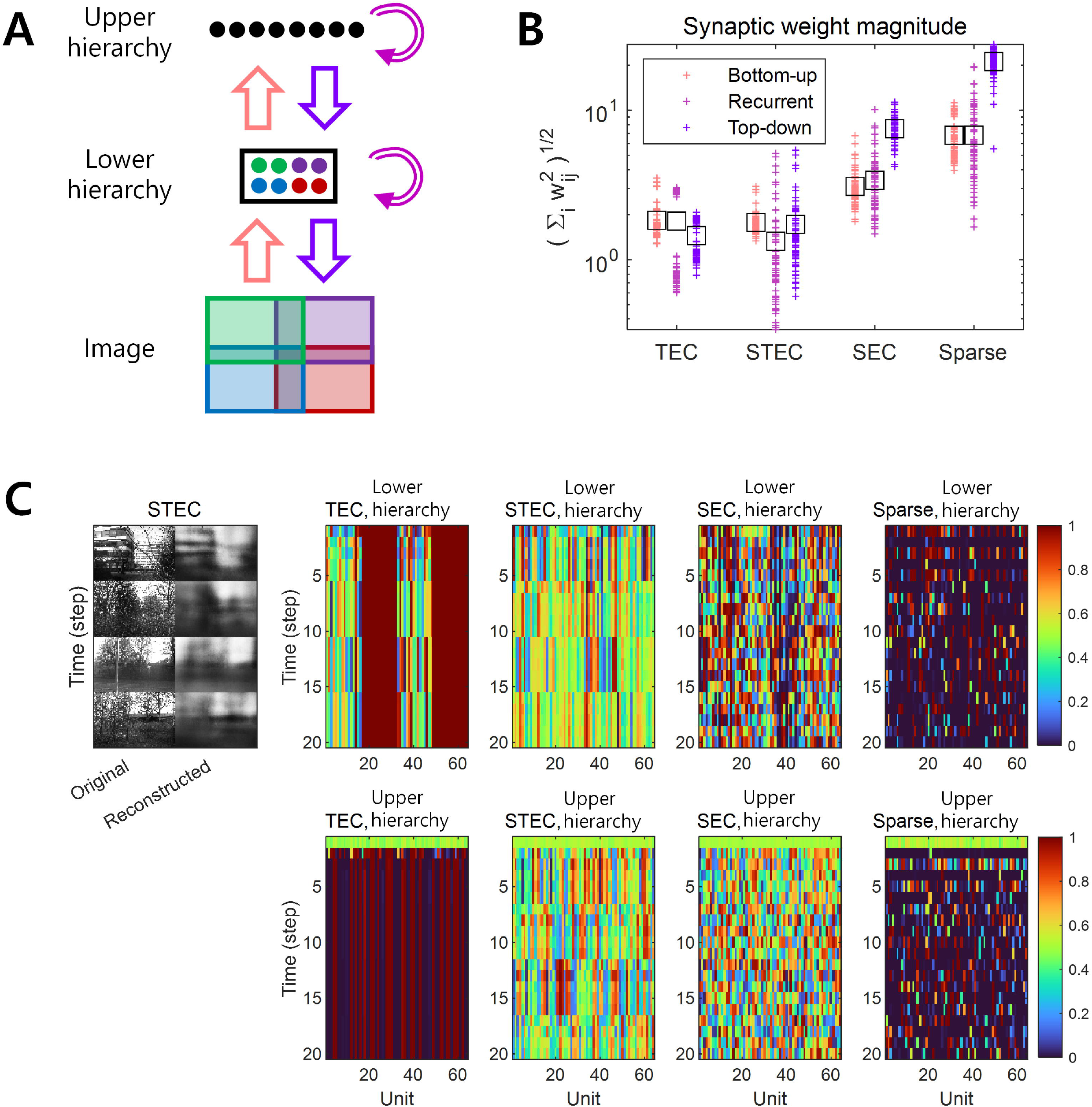
A simulation of spatio-temporally efficient coding. (A) Architecture of the simulation model. An input image is divided into four subspaces with overlaps, denoted by different colours. A subset of neuronal units in lower hierarchy receive corresponding parts of the image (matching colours). There is no spatial correspondence between lower and upper hierarchies. Pink, magenta, and purple arrows indicate bottom-up, recurrent, and top-down information transmissions, respectively. (B) The magnitude of synaptic weights on lower hierarchy units is compared for different conditions, where STEC: the balanced condition between temporally and spatially efficient coding objectives, TEC: temporally efficient coding alone, SEC: spatially efficient coding alone which is existing efficient coding, Sparse: sparse coding. Each cross indicates the L2 norm of synaptic weights for each unit. Black squares indicate the mean. (C) Left: Four different original input images and their reconstructions from the neuronal responses of lower hierarchy. Right: Representative neural responses for different conditions. The neural response is the output of the sigmoid function and is therefore normalised to a range between 0 and 1.

**Fig 3.**
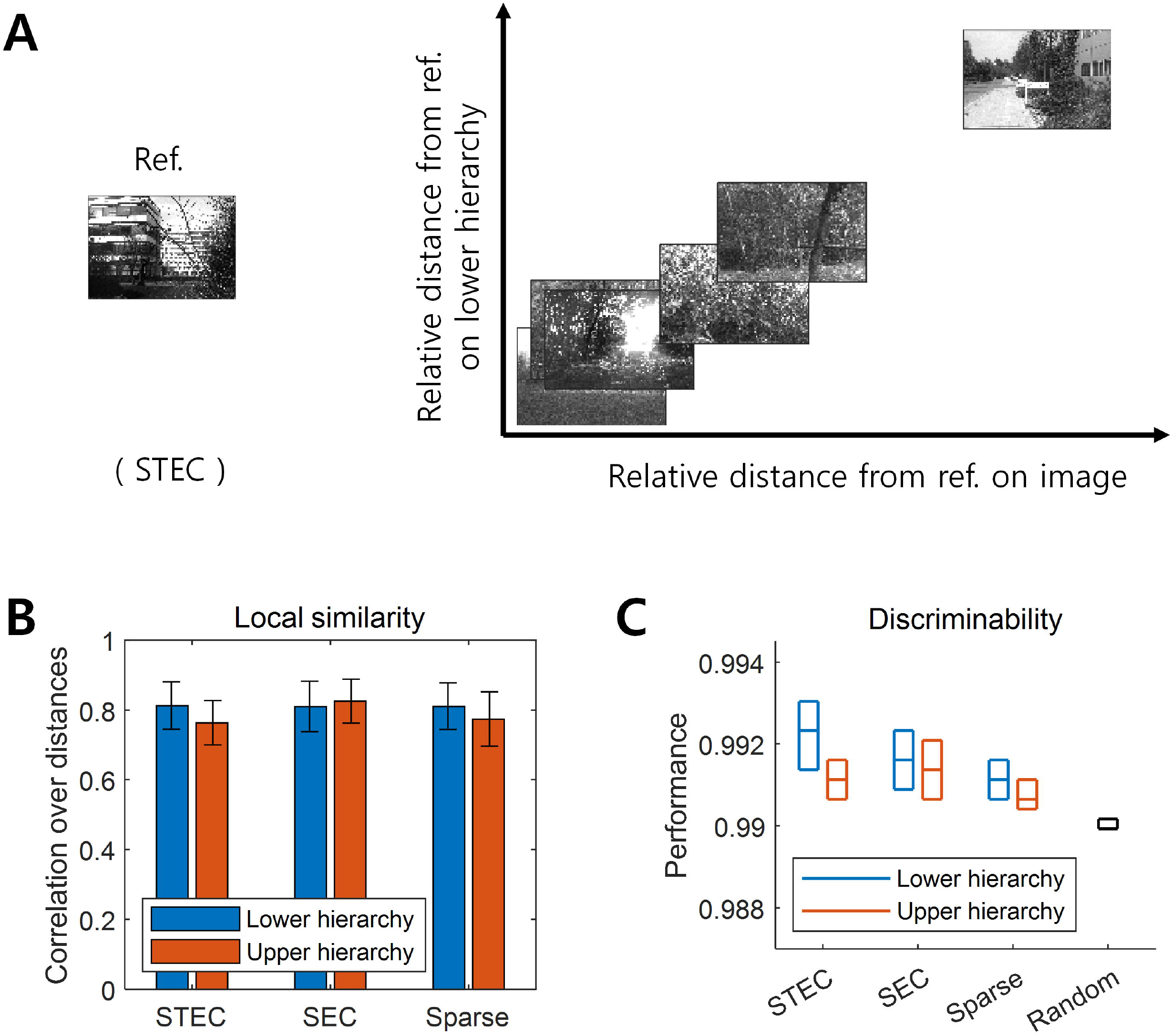
Smooth neural representations of visual images by spatio-temporally efficient coding. (A) Examples of relative Euclidean distances from the reference image (Ref. left) to other images in the natural scene image space are depicted on the horizontal axis. Relative Euclidean distances from the neural representations of the reference image at hierarchy 1 to those of corresponding other images are depicted on the vertical axis. (B) Local similarity based on correlations of distances between overall natural scene images (N = 42 out of 4212; 1%) (global feature based distance) with those of corresponding neural responses (Euclidean distance). Error bars indicate standard deviations. (C) Discriminability of neural responses for images (N = 4170 out of 4212; 99%). Three horizontal lines in each box indicates 25 %, 50 %, and 75 % levels of data, respectively.

**Fig 4.**
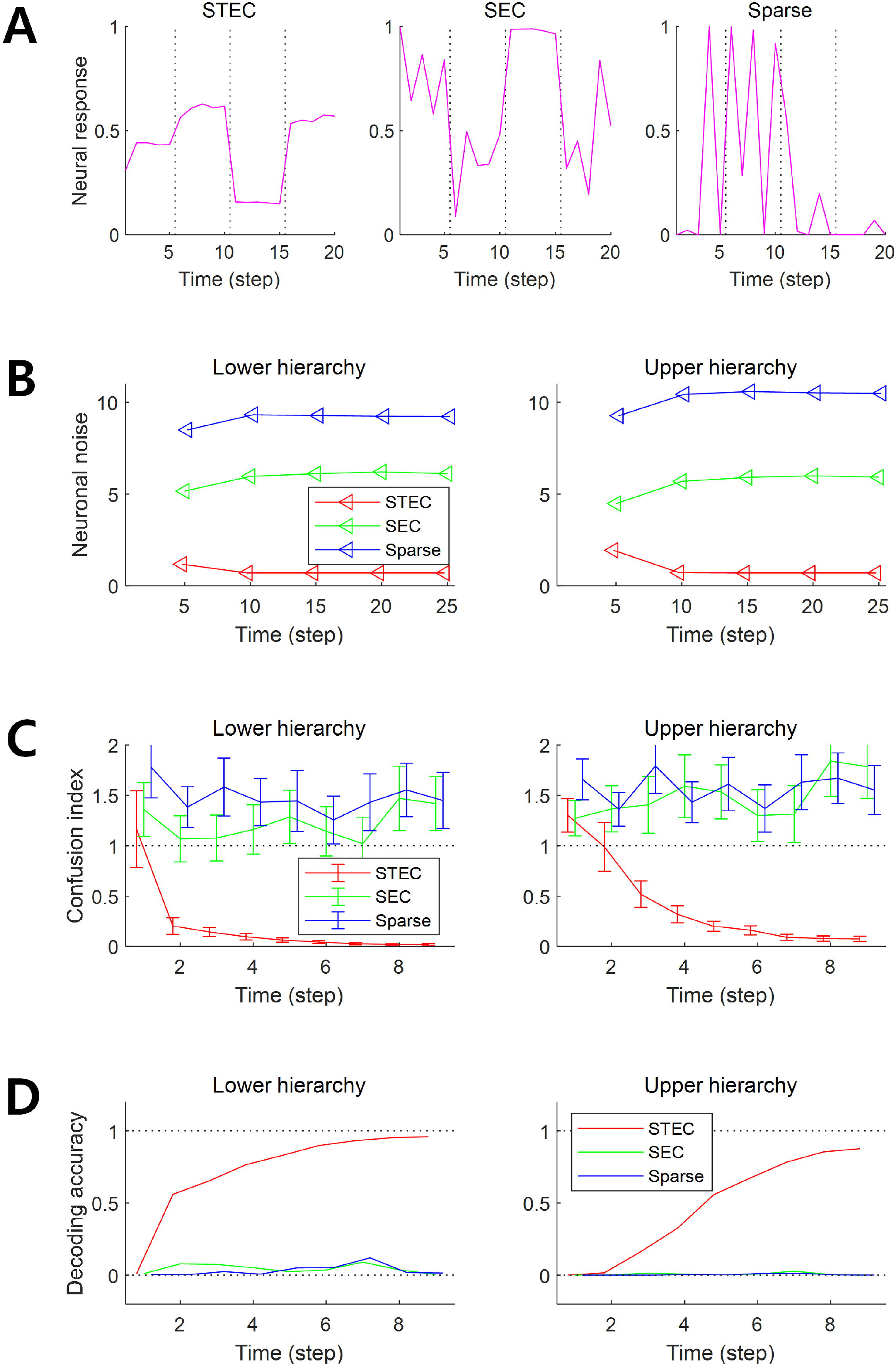
Decodable stable neural representations via spatio-temporally efficient coding. (A) Examples of neural responses in the lower hierarchy. These were neural responses of 3rd units in Fig 2C. Dotted vertical black lines indicate the presentations of new external input. (B) Neuronal noise that measured as shifted conditional entropy of neural responses given stimulus. Conditional entropy was measured by collecting neural responses at every five time step. (C) Confusion index that a measure of how much it confuses neural responses to one stimulus with neural responses to another similar stimulus. The dotted lines denote the confusion index of 1 which indicates the confusion. Error bars indicate standard deviations. (D) Decoding accuracy via the naïve Bayes classifier. The upper dotted line denotes the maximum decoding accuracy, 1. The lower dotted line denotes the chance level, 1/4212. Error bars indicate standard deviations.

## 2. Materials and Methods

### 2.1. Spatio-temporally efficient coding

Predictive coding utilizes both bottom-up and top-down pathways to describe neural representations in hierarchical structures in the brain. Nevertheless, predictive coding has several theoretical shortcomings: 1) the hierarchical structure would require an additional information processing subsystem to perform this inference to minimise the prediction error and 2) predictive coding requires the presence of error units in the hierarchical structure to represent this prediction error, yet such error units remain as hypothetical entities.

Existing efficient coding (Attneave, 1954; Barlow, 1961; Laughlin, 1981) does not require additional information processing subsystems or virtual error units, which are shortcomings of predictive coding. Unfortunately, however, existing efficient coding does not consider hierarchical structures. We, therefore, propose a novel computational principle for visual hierarchical structures as spatio-temporally efficient coding underscored by the efficient use of given resources in both neural activity space and processing time. The derivation of spatio-temporally efficient coding is as follows.

A possible approach to overcome the shortcomings of both bottom-up processing and predictive coding in visual hierarchical structures is to make bottom-up information transmissions similarly to top-down information transmissions across hierarchies, instead of transmitting bottom-up prediction errors. For example, context-independent bottom-up predictions and context-dependent top-down predictions (Teufel and Fletcher, 2020). Such bidirectional information transmissions eliminate the necessity for hypothetical error units, while presumably elucidating the neural responses of hierarchical structures underlying bottom-up feature integration and top-down predictive coding. A neural coding principle underlying bidirectional information transmissions of hierarchical structures can be found in the theory of efficient coding that draws upon the efficient use of given resources (Laughlin, 2001; Bullmore and Sporns, 2012), which crucially include limited time resources related to processing speed (Griffiths et al., 2015; Lieder and Griffiths, 2020). A possible solution to promote the most efficient use of limited time resources by the bidirectional information transmission system is to render present neural responses similar to future ones before the occurrence of future neural responses. This can be achieved by minimising the temporal differences between present and future neural responses. Accordingly, we consider this temporal difference minimisation as our *learning principle,* referred to as *temporally efficient coding.* Here, *inference* simply refers to a bidirectional information transmission mediated by top-down and bottom-up pathways. Unlike inference in predictive coding, which requires further error minimisation, inference in temporally efficient coding involves simple single-step information transmission.

Temporally efficient coding involves a trivial solution: neural responses do not change to changes in external events. This trivial solution is comparable to the dark-room problem of predictive coding or free-energy principle, where an agent stays and is unchanged in a dark room with no surprise or unpredicted parts (Friston et al., 2012; Clark, 2013). We circumvent this issue by adding a complementary neural coding (learning) principle that maximises the informational entropy of neural responses to alter neural responses to changing external events. It maximises the neural response space available to represent the external world under the constraints of both the number of neurons and maximum firing rates. Maximal entropy coding indicates that the system uses spatial resources of neural responses efficiently (Attneave, 1954; Barlow, 1961; Laughlin, 1981), referred to as *spatially efficient coding*. By combining spatially efficient coding and temporally efficient coding, we propose a neural coding principle termed *spatio-temporally efficient coding* (Fig 1).

Spatially efficient coding has the same objective as existing efficient coding (Barlow, 1961; Laughlin, 1981), which minimizes informational redundancy because it increases the difference between neural responses. Temporally efficient coding, on the other hand, can be regarded to increase informational redundancy, as it reduces the difference between neural responses. Two seemingly opposing coding objectives can be reconciled by isolating mechanisms that decrease the differences between consecutive neural responses on time domain and those that increase the differences between neural responses to the apparently different external world. In the real brain, it can be explained that different mechanisms are applied depending on the degree of difference in neural response. In the implementation of the present study, two coding objectives were applied in different ways. As an implementation of temporally efficient coding, we minimize the difference between consecutive neural responses on the time. In spatially efficient coding, the neural responses to the apparently different external stimuli are implemented in a minibatch method that simultaneously learns different images. We maximized the difference between neural responses to different images within each time step (see, Implementation of spatio-temporally efficient coding).

Temporally efficient coding trains the present neural response to be similar to the future neural response in order to efficiently use a given time resource. The temporal trajectory of the neural response is smoothed as the difference between the present and future neural responses is minimised. It thereby minimises the size of the space represented, when a single stimulus (stimulus in the external world) is represented on the time domain. In other words, it reduces neuronal noise (also see, Fig 2C) which is defined as the uncertainty of a neural response for given stimulus (Borst and Theunissen, 1999). This is to decrease the conditional entropy of neural response given stimulus, *H*(*X*|*S*) where *X* indicates neural response and *S* indicates stimulus. Spatially efficient coding increases *H*(*X*). Spatio-temporally efficient coding, thus, increases the Shannon mutual information *I*(*X*;*S*) = *H*(*X*) – *H*(*X*|*S*) simultaneously in both terms: *H*(*X*) and – *H*(*X*|*S*). This is also the definition of another existing efficient coding (Friston, 2010). Spatio-temporally efficient coding in hierarchical structures, therefore, can also be seen as an extension of efficient coding into hierarchical structures on time domain.

Neural system homeostasis is associated with maximisation of mutual information between neural responses and external stimuli (Toyoizumi et al., 2005; Sullivan and de Sa, 2006). Since spatio-temporally efficient coding increases the Shannon mutual information between neural responses and external stimuli, it is related to homeostasis. In particular, temporal difference minimisation of neural responses in temporally efficient coding is reminiscent of homeostasis of energy metabolism. Smoothing the temporal trajectory of a neural responses reduces the variance of the neural response distribution so that the neural response stays within a certain range. This is also a consequence of the homeostatic plasticity (Turrigiano and Nelson, 2004) of the brain (also see, Fig 5).

**Fig 5.**
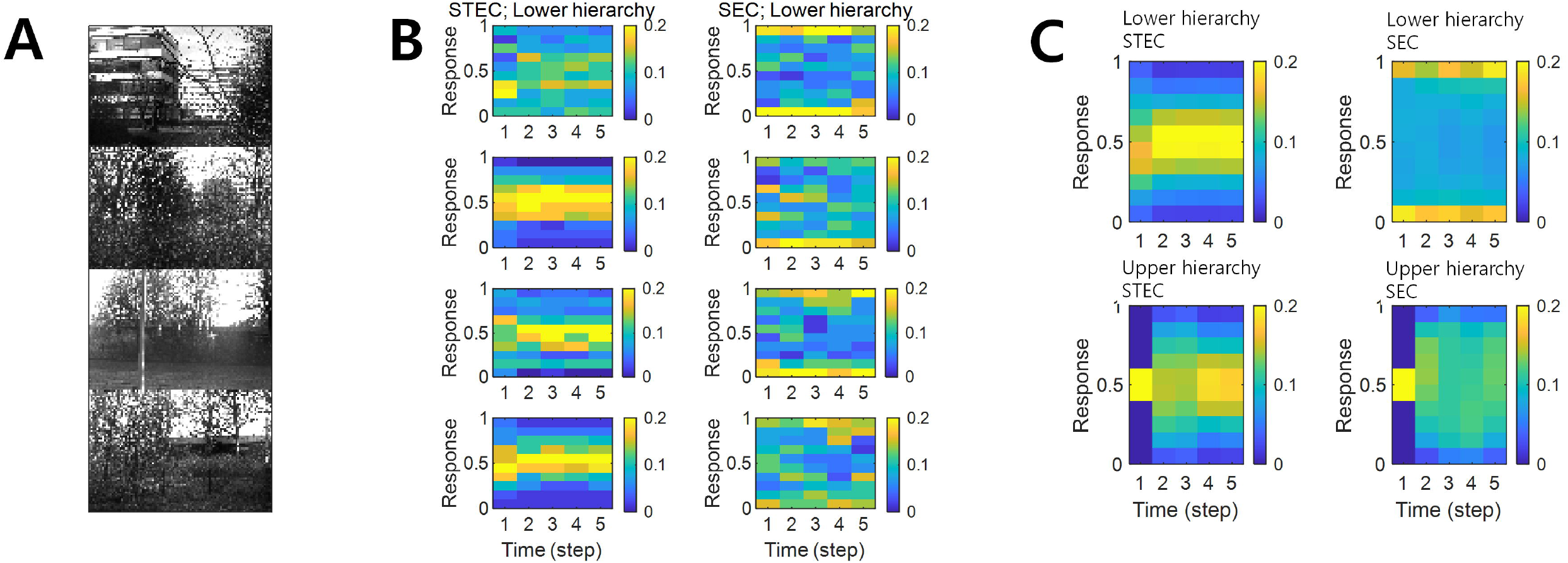
Neural response distributions. (A) Examples of natural scene images. (B) Examples of neural response distributions of lower hierarchy units for each of the five time steps of bidirectional information transmissions, corresponding to images in (A). The colour scale indicates the proportion. (C) Overall neural response distributions in response to all input images (either STEC or SEC) at lower and upper hierarchies, respectively.

As mentioned earlier, smoothing the temporal trajectories of neural responses reduces the spatial extent of neural responses to static stimuli. This leads to a rapid stabilization of the neural response to the static stimulus (also see, Fig 2C and 4). The rapid stabilization of neural responses shows the characteristic of temporally efficient coding, which makes efficient use of given time resources. Another effect of this smoothing the temporal trajectory of neural response is to render smooth neural representations that locally preserves the structure of the external world. If a stimulus is static or changes smoothly, making the temporal trajectory of the neural response smooth is to render a similar neural response to the similar stimuli. This is smooth neural representations that locally preserves the structure of the external world (also see, Fig 3).

### 2.2. Relation to other works

Existing efficient coding (Attneave, 1954; Barlow, 1961; Laughlin, 1981) is a widely accepted coding principle. However, existing efficient coding does not consider hierarchical structures. Spatio-temporally efficient coding started with a different kind of assumption from the existing efficient coding that given resources are used efficiently. However, spatio-temporally efficient coding can be seen as an extension of efficient coding to hierarchical structures. As mentioned previously, temporally efficient coding decreases the conditional entropy of neural response given stimulus, *H*(*X*|*S*) where *X* indicates neural response and S indicates stimulus. Spatially efficient coding increases *H*(*X*). Spatio-temporally efficient coding, thus, increases the Shannon mutual information *I*(*X*;*S*) = *H*(*X*) – *H*(*X*|*S*) simultaneously in both terms: *H*(*X*) and – *H*(*X*|*S*). This is also the definition of another existing efficient coding (Friston, 2010). Since spatio-temporally efficient coding can be applied to a hierarchical structure, spatio-temporally efficient coding can be viewed as an extension of efficient coding to hierarchical structures on time domain.

It is noteworthy that minimising temporal differences in upper hierarchies may, at first glance, be reminiscent of the slow feature analysis (Wiskott and Sejnowski, 2002; Berkes and Wiskott, 2005; Creutzig and Sprekeler, 2008). However, the upper hierarchies of the proposed spatio-temporally efficient coding need to change dynamically to react rapidly changing visual inputs and neural responses of the lower hierarchies, whereas the upper hierarchies of the slow feature analysis need to change slowly. Minimising temporal differences in spatio-temporally efficient coding does not aim at extracting slow features, but rather aim at the smooth representation of rapidly changing visual inputs with stabilizing neural responses as quickly as possible.

The hierarchical structure of the brain, which ascends to the upper hierarchy from the input by the bottom-up pathway and then descends to the input by the top-down pathway again, resembles the structure of autoencoders (Bourlard and Kamp, 1988; Hinton and Zemel, 1993) of artificial neural networks. Objective functions of spatio-temporally efficient coding applied to the hierarchical structure for neural representations can be viewed as a kind of objective function applied to autoencoders. Many autoencoders have also been used for representation learning (Bengio et al., 2013). Typical examples include sparse autoencoders (Ranzato et al., 2006), denoising autoencoders (Vincent et al., 2008), and contractive autoencoders (Rifai et al., 2011). However, these examples do not take into account the passage of time like spatio-temporally efficient coding. Similar to slow feature analysis, there have been studies on the properties of cells in the visual cortex using temporal coherence to obtain slow representations (Hurri and Hyvärinen, 2002; Zou et al., 2011). In particular, one of these studies has been conducted using models that partially include autoencoders (Zou et al., 2011). However it is different from spatio-temporally efficient coding in that it lacks an aspect of dynamically reacting to changes in external input or neural responses of other hierarchies, and only partially includes autoencoders.

### 2.3. Implementation of spatio-temporally efficient coding

In the present study, visual information processing in hierarchical structures was established as biologically inspired temporal processing. Specifically, visual information processing is described as a function *f_t_* for both image *X_image_* and neural responses *X_h_* in each hierarchy *h* such that it maps from *X_image_* and *X_h_* at time *t* – 1 to those at time *t*:

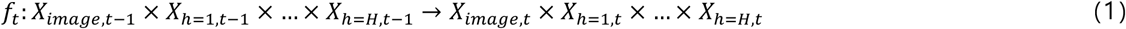

where *X*_*h,t*-1_ and *X_h,t_* are the neural responses *X_h_* at time *t* – 1 and *t*, respectively, at hierarchy *h*. For convenience, *X*_*h*=0_ denotes *X_image_*, in particular *X*_*h*=0,*t*_ denotes the image presented at time *t*. The details of the visual information processing *f_t_* are as follows: If *h* > 0, then

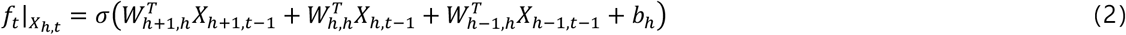

where *X_h,t_* is an *X_h_* value vector at time *t, f_t_*(·)|_*X_h,t_*_ indicates restricting the range of the function value *f_t_*(·) to *X_h,t_*, *W*_*h*+1,*h*_ is a synaptic weight matrix from hierarchy *h* + 1 to *h, T* is the transpose of a matrix, *b_h_* is a bias vector at hierarchy *h*, and *σ*(·) is a sigmoid function. The terms of 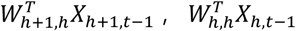, and 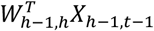 indicate top-down, recurrent, and bottom-up information transmissions, respectively. In case of *h* + 1 > *H* the 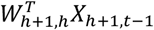 term would be omitted. If *h* = 0, then

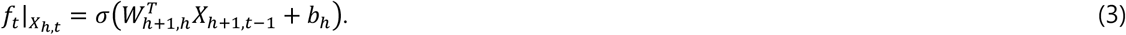

If *h* > 0, *X_h,t_* = *f_t_*|*x_h,t_*. On the other hand, when *h* = 0, *X_h,t_* is the image presented at time *t*, not *f_t_*|*X_h,t_*. This function *f_t_* makes *inferences* using spatio-temporally efficient coding. Unlike inference in predictive coding (Rao and Ballard, 1999; Spratling, 2017) that requires additional processes such as minimisation of prediction errors, inference in spatio-temporally efficient coding is a function value *f_t_* itself.

*Learning* in spatio-temporally efficient coding minimises the ensuing objectives of both temporally and spatially efficient coding. The objective of temporally efficient coding is given by:

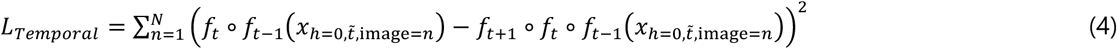

where ° is the function composition, 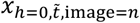 indicates that the image is fixed to the *n*th sample throughout the temporal processing (i.e., *X*_*h*=0,*t*–2_ = *X*_*h*=0,*t*–1_ = *X*_*h*=0,*t*_ is *n*th image sample), squaring is operated component-wise. In case of *h* = 0, 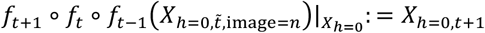 which is the image presented at time *t* + 1 while 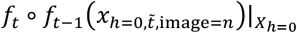 is the function value as inference. The function composition *f_t_* ° *f*_*t*–1_ of two functions (inferences) in the *L_Temporal_* allows the simultaneous learning of all information transmissions across all hierarchies because the depth of hierarchy is *H* = 2 in the present study (bottom-up, recurrent, and top-down information transmission across the hierarchy of depth *H* = 2). The minimisation of the given objective minimises the temporal differences between present and future neural responses. The objective of temporally efficient coding is to render present neural responses similar to future neural responses that has not yet arrived. By doing so, it uses the given time resources efficiently. This minimisation of temporal difference creates a learning effect in which the temporal trajectory of the neural response becomes smooth. The expected effect of this learning effect is to quickly stabilize neural responses when a static external stimulus is given (also see, Fig 2C and 4). In that aspect, it is also to use the given time resources efficiently.

The objective of spatially efficient coding is given by

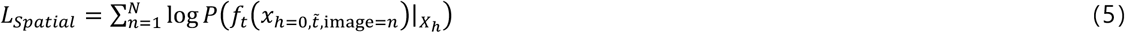

Where *h* > 0 in *X_h_* (so, 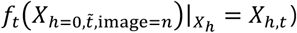 and *P*(·) is a probability. Because the term *L_Spatial_* is an estimation of negative informational entropy, minimising the objective maximises the informational entropy of *f_t_*. This objective increases the entropy of each neuron, and it can be seen that an increase in marginal entropy in each neuron will increase the joint entropy of the entire system. Therefore, spatially efficient coding maximises the entropy of individual neurons as in Laughlin’s study (Laughlin, 1981) and consequently the entropy of the entire system. The objective of spatially efficient coding is to render different neural responses to different inputs from the external world. We, therefore, can overcome the trivial solution of temporally efficient coding, where there is no change in neural response despite changes in the external world. This thereby does not conflict with the learning of temporally efficient coding, which decreases differences in consecutive neural responses in the time domain to inputs of one stream.

To minimise *L_Spatial_*, which is the objective of spatially efficient coding, it is necessary to calculate the probability *P*(·) in *L_Spatial_* (equation 5). Instead of calculating the exact probabilities, we obtained pre-normalised densities in the sense of probabilities without a partition function. As the value of the partition function is fixed, it does not affect the minimisation process. Note that, when *h* > 0, *X_h,t_* is the space of all possible neural responses, i.e., *X_h,t_* = [0,1]^dim *X_h,t_*^. Let *x* ∈ *X_h,t_* be a value of 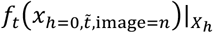. Kernel density estimation was used to obtain *P*(*x*). Using a Gaussian kernel with width 0.1(dim *X_h,t_*)^1/2^, the neural response density *Q*(*x*) and compensation density *Q*′(*x*) at *x* ∈ *X_h,t_* were obtained. Then, the pre-normalised density of interest is

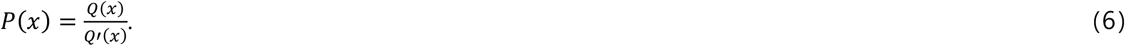

The neural response density *Q*(*x*) is obtained by kernel density estimation of neural responses on *X_h,t_*. The compensation density *Q*′(*x*) is obtained using pseudo-uniformly generated samples on *X_h,t_* instead of the neural responses. (For sparseness constraint, i.e., sparse neural responses, *Q*′(*x*) is obtained using pseudo-uniformly generated samples on [-1,1]^dim *X_h,t_*^ instead of *X_h,t_* = [0,1]^dim *X_h,t_*^, so that *Q*′(*x*) has a fat distribution around zero. See, Fig 6) The compensation density is necessary to compensate for the non-uniform intrinsic expectation of *Q*(·) resulting from the fact that *X_h,t_* is bounded. At the boundary of *X_h,t_*, the density of neural responses, *Q*(*x*), measured by kernel density estimation is decrease. This intrinsic decrease corresponds to *Q*′(*x*). We compensated for the decrease by dividing *Q*(*x*) by *Q*′(*x*).

**Fig 6.**
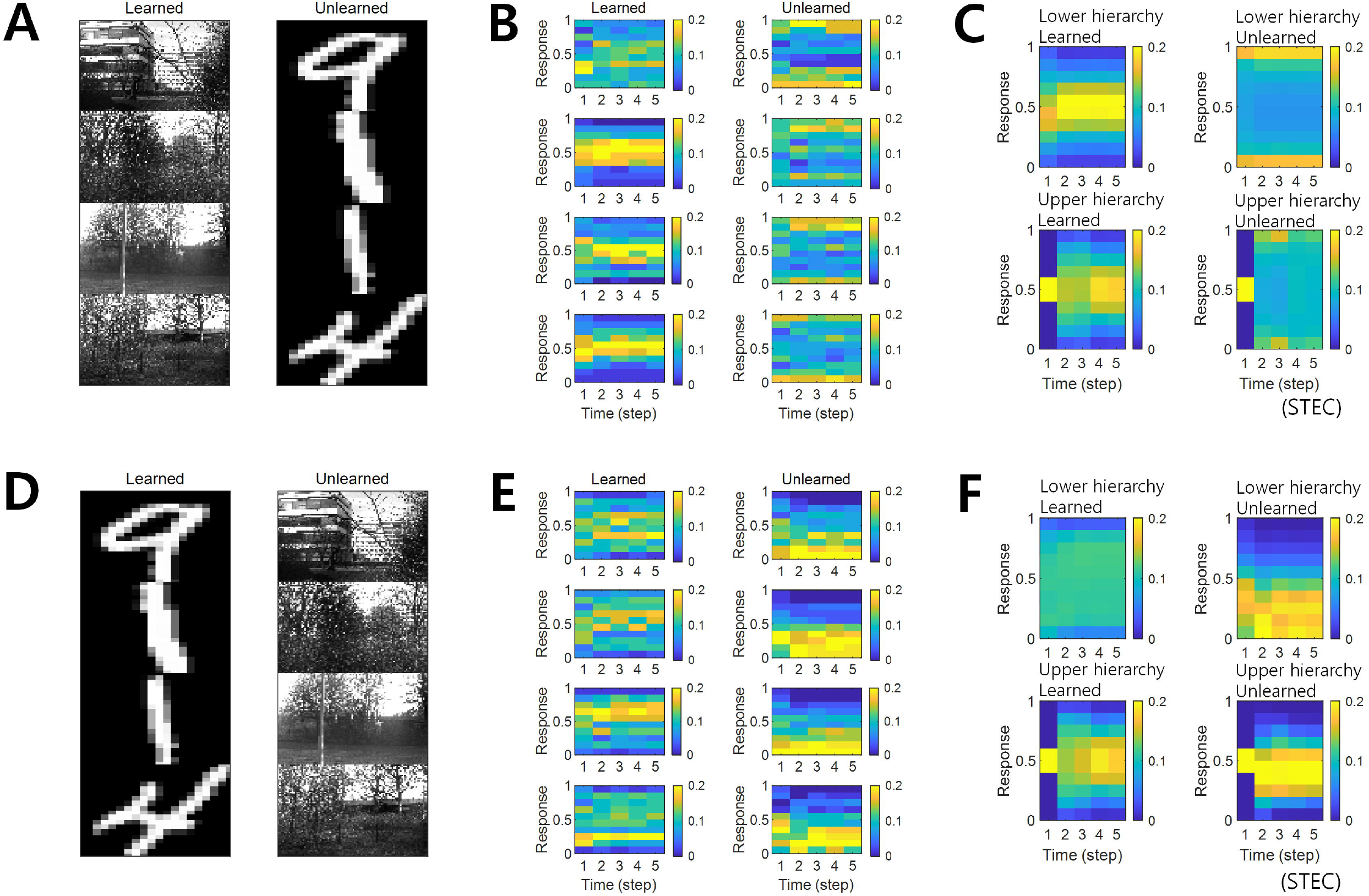
Neural response distributions for learned and unlearned inputs. (A) Natural scene images are used as learned inputs (for learning in the visual hierarchical structure) and handwritten digit images are used as unlearned inputs (not used in learning). (B) Examples of neural response distributions of lower hierarchy units for each of the five time steps of bidirectional information transmissions, corresponding to images in (A). The colour scale indicates the proportion. (C) Overall neural response distributions in response to all input images (either learned or unlearned) at lower and upper hierarchies, respectively. (D), (E), (F) are similar to (A), (B), (C), but handwritten digit images are used as learned inputs and natural scene images are used as unlearned inputs.

Finally, the objective of spatio-temporally efficient coding is a linear combination of those two objectives:

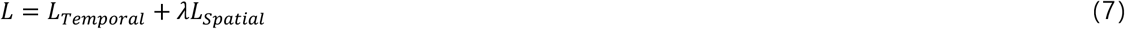

where *λ* is a regularisation parameter. A smaller *λ* indicates a greater emphasis on the temporally efficient coding objective, whereas a larger *λ* indicates the opposite. As mentioned previously, temporally efficient coding decreases the conditional entropy of neural response given stimulus, *H*(*X*|*S*) where *X* indicates neural response and *S* indicates stimulus. Spatially efficient coding increases *H*(*X*). Hence, the regularisation parameter *λ* can be seen as controlling the balance between *H*(*X*) and – *H*(*X*|*S*). It is a modification of fixed balance of Shannon mutual information *I*(*X*;*S*) = *H*(*X*) – *H*(*X*|*S*) of the existing efficient coding (Friston, 2010).

Temporal trajectories of neural responses are smoothed by temporally efficient coding, but this does not mean just slow neural representations. By spatially efficient coding, different neural responses to different inputs from the external world should be exhibited. Therefore, it should show rapid changes in neural responses to sudden changes in the external world (fast representation). This is the difference from slow feature analysis (Wiskott and Sejnowski, 2002; Berkes and Wiskott, 2005; Creutzig and Sprekeler, 2008) or temporal coherence (Hurri and Hyvärinen, 2002; Zou et al., 2011), which targets slow neural representations. On the one hand, with temporally efficient coding, changes in neural responses should be smoothed out quickly when the external input is not changing (fast stabilization).

Suppose that Gaussian noise is added to a series of temporally correlated external inputs (e.g., static images or smoothly moved images + Gaussian noise). If these noisy external inputs are still temporally correlated, then smoothing the temporal trajectory of neural responses (by temporally efficient coding objective *L_Temporal_)* is rendering temporally similar neural responses to temporally similar inputs. It is also smooth neural representations that locally preserves the structure of the external world. Moreover, if the noise is provided independently of the input, the effect of the noise on neural representations will be dispelled by multiple independent trials. Therefore, temporally efficient coding objective *L_Temporal_* increases the fidelity of the neural representation with respect to a Gaussian noise.

Smooth neural representation also means making different neural responses to different inputs. In other words, it increases discriminability for different inputs. This is achieved by spatially efficient coding objective *L_Spatial_* that maximises the entropy of neural responses. Assumed that Gaussian noise is added to external inputs. Let *D*_true_ be a binary-valued random variable for the discrimination between two actually different inputs such that *D*_true_ = 1 means the discrimination that two noised inputs differ and *D*_true_ = 0 means the discrimination that two noised inputs are same. *D*_true_ is probabilistic because of the Gaussian noise mentioned earlier. Let *D*_response_ be a binary-valued random variable for the discrimination between two neural responses for inputs such that *D*_response_ = 1 means the discrimination that two neural responses differ and *D*_response_ = 0 means the discrimination that two neural responses are same. Let *P*_true_ and *P*_response_ be the probability mass functions of *D*_true_ and *D*_response_, respectively. Spatially efficient coding objective *L_Spatial_* decreases the Kullback-Leibler divergence from response to *P*_response_, i.e., *P*_true_||*P*_response_). Hence it also decreases the binary cross entropy *H*(*P*_true_) + *D*_KL_(*P*_true_||*P*_response_) where *H*(·) is an informational entropy.

In the present study, the depth *H* of hierarchies was set to 2, the minimum depth to realise both bottom-up and top-down pathways in the same hierarchy. Images with 64 × 96 size were divided into four overlapping 38 × 58 patches. Each patch was connected to 16 of 64 lower hierarchy units. Further, 64 lower hierarchy units were fully connected to 64 upper hierarchy units. All units in each hierarchy are fully connected to each other (Fig 2A).

Because *L* = *L_Temporal_* + *λL_Spatial_* is differentiable, the minimisation of the objective in spatio-temporally efficient coding was performed with a gradient descent. The Adam optimiser (Kingma and Ba, 2015) was used to perform the stochastic gradient descent with momentum. The parameters of the Adam optimiser used in this study were *α* = 0.001, *β*_1_ = 0.9, *β*_2_ = 0.999, and *ϵ* = 10^-8^. The optimisation lasted 10^4^ iterations for each repetition and restarted with five repetitions. For each iteration, the duration of temporal processing *f_t_* was five (i.e., *t* ∈ [1,5]), and the minibatch size was 40. For a given image, the duration of temporal processing of *f_t_* was given as five (i.e., *t* ∈ [1,5]) in both learning and inference, because four time steps are required for the image information to reach the top hierarchy and return over *H* = 2 hierarchies, in addition to one time step to obtain future neural responses. In learning, after temporal processing of five durations was finished, new temporal processing begins, and the initial values of neural responses of new temporal processing were the last neural responses values of the previous temporal processing.In our simulations, we repeatedly exposed the hierarchical structure to natural scene images, which enabled it to learn the bidirectional information transmissions between top-down and bottom-up hierarchies using spatio-temporally efficient coding with a range of the balancing parameter *λ.* Successful learning was confirmed by minimising or stabilising *L* during learning. Further, we verified that the learned hierarchical structure could successfully reconstruct an input image, as shown in Fig 2C.

Simulation codes for spatio-temporally efficient coding are available from https://github.com/DuhoSihn/Spatio-temporally-efficient-coding (Sihn, 2021).

### 2.4. Datasets

For the simulations, van Hateren’s natural scene image dataset (van Hateren and van der Schaaf, 1998) was used. The dataset was downloaded from http://bethgelab.org/datasets/vanhateren/. The images were downsized to 64 × 96 pixels. For the comparison tests, the MNIST handwritten digit dataset (Lecun et al., 1998) was used. The dataset was downloaded from http://yann.lecun.com/exdb/mnist/. The images were resized to 64 × 96 pixels to fit the images used in the simulations. All image data were rescaled between 0 and 1.

### 2.5. Computation of receptive fields

Images of 64 × 96 size with randomly selected 16 × 16 subregions were created to compute the receptive field. These 64 × 96 images had a value of one in the 16 × 16 subregions and zero otherwise. Receptive fields were calculated based on the average neural response when the visual hierarchy (*X*_*h*=1_ ×⋯×*X_h=H_*) was exposed to these images. Specifically, the value of the receptive field at a pixel was defined as the average neural response to an image with a value of one at this pixel. This averaging method was inspired by the previous study in case of binary neural responses (firing or not) (Ohzawa et al., 1996). The difference is that the previous study was for image averaging and the present study was for neural response averaging.

## 3. Results

### 3.1. Balancing between temporally and spatially efficient coding

The temporally efficient coding objective that minimises the temporal difference between present and future neural responses smooths the temporal trajectory of neural responses. The spatially efficient coding objective that maximises the informational entropy of neural responses renders different neural responses to different inputs. We can predict that when the temporally efficient coding objective and the spatially efficient coding objective are properly balanced, different neural responses to different stimuli and temporally smooth neural responses to the same stimuli can be expected. This means that when the external input changes, the neural response changes quickly, and when the external input does not change, the neural response quickly stabilizes. If the temporally efficient coding objective is overweighted, it can be predicted to arrive at a trivial solution in which the neural response does not change despite changes in external input. Conversely, if the spatially efficient coding objective is overweighted, it can be predicted that the temporal trajectory of the neural response is not smooth, resulting in neuronal noise. Three regularisation parameters *λ* ∈ {0,10,1000} were used to confirm these predictions through simulations. *λ* = 10 represents a balanced condition between the temporally efficient coding objective and the spatially efficient coding objective (STEC condition). *λ* = 0 represents an overweighted condition for the temporally efficient coding objective (TEC condition). *λ* = 1000 represents an overweighted condition for the spatially efficient coding objective (SEC condition). This control condition (SEC condition) is a condition in which only the spatially efficient coding objective is considered, and represents the existing efficient coding (Barlow, 1961; Laughlin, 1981). Sparse coding (Olshausen and Field, 1996; Olshausen and Field, 1997) was chosen to compare with other neural coding principles where temporal factors were not considered. The sparseness constraint (see, Implementation of spatio-temporally efficient coding) is applied to *λ* = 1000 to implement sparse coding (Sparse condition).

Based on the simulation, we confirmed this phenomenon by observing that, under STEC condition (*λ* = 10), when the external input changes, the neural response changes rapidly, and when the external input does not change, the neural response quickly stabilizes. In addition, we confirmed that under TEC condition (Λ = 0, control), we arrive at the trivial solution in which the neural response does not change even when the external input changes. We also confirmed that under SEC condition (Λ = 1000) and Sparse condition (Λ = 1000 + sparseness constraint), the temporal trajectory of the neural response is not smooth, resulting in neuronal noise (Fig 2C). This also shows the problem of applying existing efficient coding (Barlow, 1961; Laughlin, 1981) and sparse coding (Olshausen and Field, 1996; Olshausen and Field, 1997) directly to the hierarchical structure on time domain.

A change in balance between the two objectives also altered the relative strengths between bottom-up and top-down synaptic connections. Simulation results demonstrated that top-down synaptic strengths from upper to lower hierarchy were increased compared to bottom-up synaptic strengths from images to lower hierarchy as *λ* increased, with an emphasis on spatially efficient coding (Fig 2B, see Materials and Methods section for details on the computation of synaptic strength). This finding suggests that the balance between spatially and temporally efficient coding could be related to the balance between bottom-up and top-down synaptic strengths in the brain.

### 3.2. Smooth neural representations of the external world

As a major function of the brain is to represent the external world (deCharms and Zador, 2000; Kriegeskorte and Diedrichsen, 2019), we investigated whether spatio-temporally efficient coding creates appropriate neural representations of the external world. Spatio-temporally efficient coding smooths the temporal trajectory of neural responses while having different neural responses to different external inputs. This effect renders smooth neural representations that locally preserves the structure of the external world. We quantified the smoothness of neural representations by examining the correlations between a natural scene image space and a neural response space. Specifically, we first measured the global feature based distance (Di Gesù and Starovoitov, 1999) from a natural scene image (i.e., reference image) to other natural scene images in the image space and the Euclidean distances from the neural responses for the reference image to those for other compared images in the neural response space (Fig 3A). It has been proven that global feature based distance reflects differences in images in a way that humans actually perceive (Di Gesù and Starovoitov, 1999). When calculating the distances between neural responses, we used the neural responses at five time steps after receiving a given image based on the assumption that the neural responses became temporally steady after five steps of temporal processing (see Fig 1B). We then measured the Pearson’s linear correlation between the distances in the image space and those in the neural response space. A high correlation indicates that neural responses tend to be similar when the visual system perceives natural scenes close to each other (in the sense of global feature based distance). We set each image sample as a reference and repeated the calculations for the correlation across all image samples. We defined local similarity as the average correlation obtained by using the distances from one image to its neighbouring images only (1% of all images). We also measured discriminability as the proportion of dissimilar neural responses among dissimilar images. This dissimilarity was defined as 99% of all, based on certain distances (the global feature based distance for images, the Euclidean distance for neural responses). A high discriminability indicates that if the image is different, the neural response will also be different.

If the learned neural representations are smooth, i.e. locally preserving the structure of the external world, both local similarity and discriminability will be high. We can predict that when the temporally efficient coding objective and spatially efficient coding objective are balanced (STEC condition), neural representations are smooth, so that local similarity and discriminability are high. From the simulations, we confirmed that local similarity and discriminability do not lower in STEC condition than SEC or Sparse condition (Fig 3B and 3C).

### 3.3. Decodable stable neural representations

In a hierarchical structure in which information is exchanged in both directions, if the information represented by the upper and lower hierarchies at the same time is different, it is difficult to obtain a stable neural response on time domain for an external input. The reason is that if the information represented by the upper and lower hierarchy are different, different information is exchanged, and thus the information represented next time may be also different. These inter-connected structures could also produce chaotic dynamics (Rubinov et al., 2009; Tomov et al., 2014).

The objective of temporally efficient coding is to render present neural responses similar to future neural responses that has not yet arrived. This minimisation of temporal difference creates a learning effect in which the temporal trajectory of the neural response becomes smooth. It thereby minimises the size of the space represented, when a single stimulus is represented on the time domain. In other words, it reduces neuronal noise which is defined as the uncertainty of a neural response for given stimulus (Borst and Theunissen, 1999). The expected effect of this learning effect is to quickly stabilize neural responses when a static external stimulus is given. The objective of spatially efficient coding is to render different neural responses to different stimuli. The expected effect of spatio-temporally efficient coding is to render decodable stable neural representations which is an appropriate function to hierarchical brain structures on the time domain.

As a result of the simulation, different neural responses to different stimuli were shown in the STEC condition, and these neural responses were rapidly stabilized. On the other hand, neural responses were less stabilized under SEC and sparse conditions (Fig 4A). The factor affecting the stabilization of the neural response was the amount of neuronal noise. To quantify the amount of neural noise, conditional entropy of neural responses given stimulus was measured. Conditional entropy was measured by collecting neural responses at every five time step. Probability was estimated by kernel density estimation without a partition function, and thus shifted conditional entropy was measured. The STEC condition showed lower neuronal noise than the SEC and Sparse conditions (Fig 4B).

We measured how much this neuronal noise makes to confuse the neural response to one stimulus with the neural response to another similar stimulus. This is represented by the confusion index. Here, we set the time step to 10 and measured the variability in neural responses from time step 1 (i.e., when receiving an input image) to time step 9 until convergence. Temporal variations were computed using the Euclidean distance from the neural responses at each time step (1, …, 9) to neural responses at time step 10. We also computed the Euclidean distance from the neural responses at time step 10 and those for an image that was the nearest to the given image in the image space. We compared the temporal variations above with this distance to the nearest neighbouring image by calculating the ratio of the temporal variations over the distance to the nearest neighbouring image. This ratio was called the confusion index. The confusion index exceeds 1 indicates that the noise could significantly interfere with neural representations to be confused with nearby images. Under STEC condition, the confusion index was kept below 1 on average for the majority of time steps at each hierarchy (Fig 4C). This means that the neural representations are rapidly stabilized in STEC condition. On the other hand, the confusion index was not sufficiently reduced in SEC and Sparse condition (Fig 4C). This means that sufficient stabilization of neural representations is difficult with only existing efficient coding or sparse coding.

To measure how decodable the neural response is, the neural responses of time steps 1, …, 9 were decoded using the neural responses of time steps 10 and 11 as a training set. The naïve Bayes classifier (Hastie et al., 2009) was selected as the decoder, and each of the 4212 natural scene images was defined as one class. In the STEC condition, the decoding accuracy gradually increased over time, suggesting decodable stable neural representations. On the other hand, the decoding accuracy was low in SEC and Sparse conditions.

### 3.4. Relation to homeostasis

Neural system homeostasis is associated with maximisation of mutual information between neural responses and external stimuli (Toyoizumi et al., 2005; Sullivan and de Sa, 2006). Since spatio-temporally efficient coding increases the Shannon mutual information between neural responses and external stimuli, it is related to homeostasis. In particular, temporal difference minimisation of neural responses in temporally efficient coding is reminiscent of homeostasis of energy metabolism. Smoothing the temporal trajectory of a neural responses reduces the variance of the neural response distribution so that the neural response stays within a certain range. We confirmed this through simulation. In STEC condition emphasizing temporally efficient coding objective, neural responses were concentrated at the middle value than in SEC condition (Fig 5). This is also a consequence of the homeostatic plasticity (Turrigiano and Nelson, 2004) of the brain.

### 3.5. Deviant neural responses to unlearned inputs

The visual system often responds selectively to sensory inputs (Margoliash, 1983; Waydo et al., 2006). Even for the type of sensory inputs to which the visual system is responsive, unfamiliar inputs induce larger neural responses compared to familiar inputs (Huang et al., 2018; Issa et al., 2018). These large neural responses to unfamiliar inputs are thought to be due to prediction errors (Issa et al., 2018). On the one hand, spatio-temporally efficient coding renders smooth neural representations, i.e., locally preserving the structure of the external world. Because unlearned inputs differ from learned inputs, if their neural representations are smooth, their neural representations will also differ. These are deviant neural responses to unlearned inputs. Accordingly, we can expect that spatio-temporally efficient coding could predict the phenomenon of deviant neural responses to unlearned inputs without the introduction of prediction error responses mediated by error units.

In the simulations, the visual hierarchical structure learned to be learned with natural scene images, and novel handwritten digit images were used as unlearned visual inputs (Fig 6A). The simulation results revealed that neural responses were distributed over middle values for learned images and over smaller or larger values for unlearned inputs (Fig 6B and 6C), suggesting that spatio-temporally efficient coding could predict the phenomenon of deviant neural responses to unlearned inputs. The same conclusion could also be reached if the learned images were handwritten digit images and the unlearned images were natural scene images (Fig 6D, 6E, and 6F).

Simulations in the present study demonstrated that neural responses were distributed around smaller or larger extremes for unlearned inputs and around intermediate values for learned inputs (Fig 6B and 6C). In a separate analysis, we allowed the neural responses to learned inputs be distributed only around lower values (see, Implementation of spatio-temporally efficient coding, *λ* = 5), similar to sparse coding (Olshausen and Field, 1996; Olshausen and Field, 1997), and observed that neural responses of the lower hierarchy to unlearned inputs exhibited higher values (Fig 7). Under this sparseness constraint, when an external stimulus was first given, the neural response surged from zero to a large value and then stabilized again (Fig 7C and 7D).

**Fig 7.**
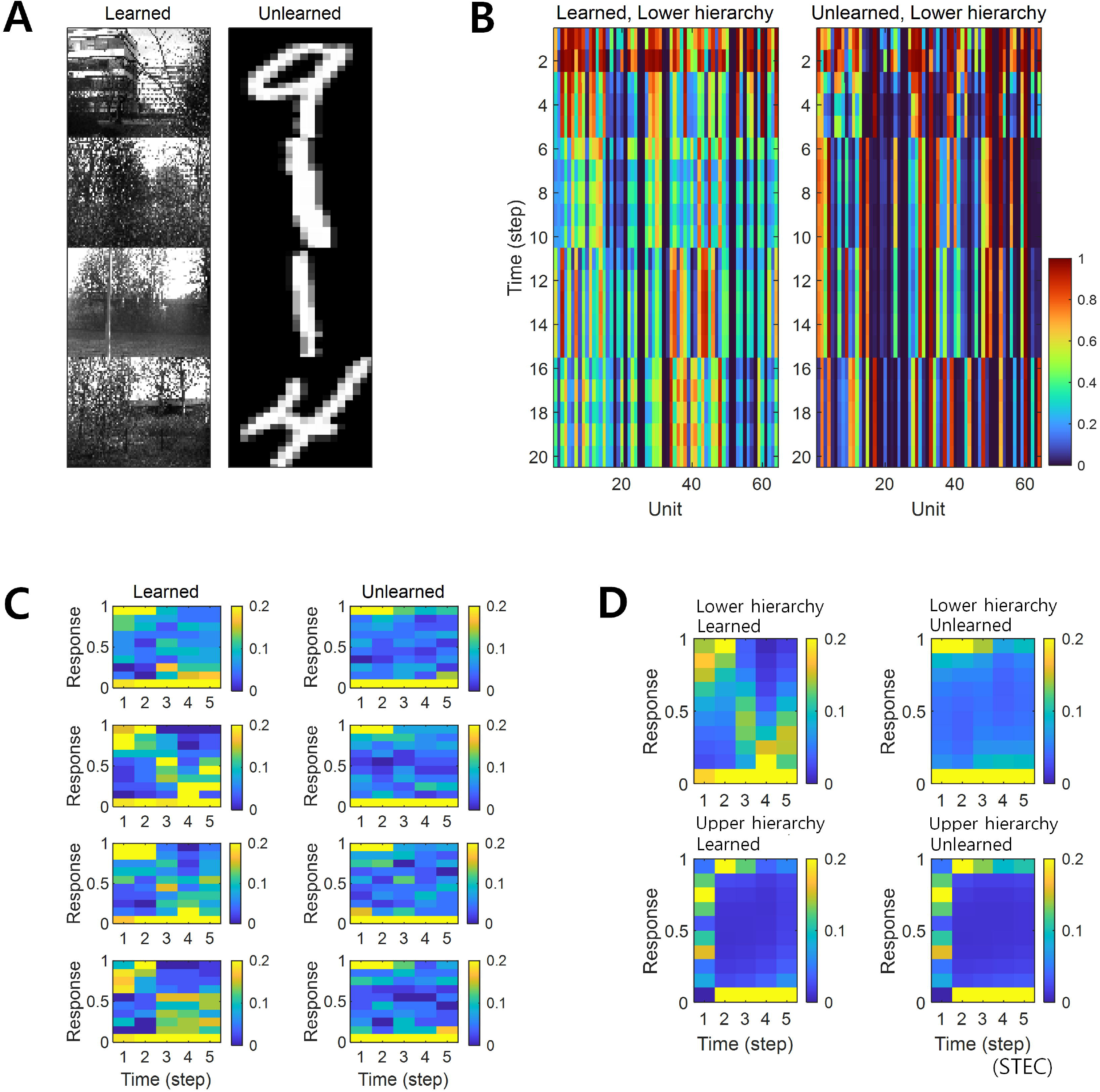
Neural response distributions for learned and unlearned inputs under sparseness constraint. (A) Natural scene images are used as learned inputs (for learning in the visual hierarchical structure) and handwritten digit images are used as unlearned inputs (not used in learning). (B) Lower hierarchy neural responses for consecutive external inputs in (A). The neural response is the output of the sigmoid function and is therefore normalised to a range between 0 and 1. (C) Examples of neural response distributions of lower hierarchy units for each of the five time steps of bidirectional information transmissions, corresponding to images in (A). Unlike (B), it is a neural response when images are presented separately rather than consecutively. The colour scale indicates the proportion. (D) Overall neural response distributions in response to all input images (either learned or unlearned) at lower and upper hierarchies, respectively.

### 3.6. Preferred orientation biases of receptive fields

Neurons in the visual system prefer horizontal and vertical orientations over oblique orientations (Furmanski and Engel, 2000; Li et al., 2003). Indeed, orientation discrimination is more sensitive to horizontal and vertical orientations than to oblique orientations (Girshick et al., 2011). This is a bias towards cardinals (Girshick et al., 2011). Since smooth neural representations of spatio-temporally efficient coding reflects the structure of the external world well, we predicted that it would also reflect the environmental statistics of natural scenes. We investigated whether units in the visual hierarchical structure that learned by spatio-temporally efficient coding of natural scene images exhibited such biases. Units in the lower hierarchy had the Gabor-like visual receptive fields, while units in the upper hierarchy had more complex visual receptive fields (Fig 8A) (also see, Computation of receptive fields in Materials and Methods section). Because the units had an oriented Gabor-like receptive field, we used oriented bar stimuli to measure the unit’s preferred orientation. We presented a static bar oriented in one of eight angles that moved in the direction perpendicular to the orientation angle (Fig 8B) and defined the response of each unit to that orientation by the largest response during presentation. We then defined the preferred orientation of each unit as the orientation that elicited the largest response. Example neural responses were shown in Fig 8C. Simulation results revealed that units clearly prefer horizontal and vertical orientations over oblique orientations (Fig 8D), consistent with the orientation bias of visual cortical neurons and in accordance with context-independent bottom-up prediction (Teufel and Fletcher, 2020). This means that smooth neural representations when well-balanced between temporally efficient coding objective and spatially efficient coding objective reflect the environmental statistics of natural scenes.

**Fig 8.**
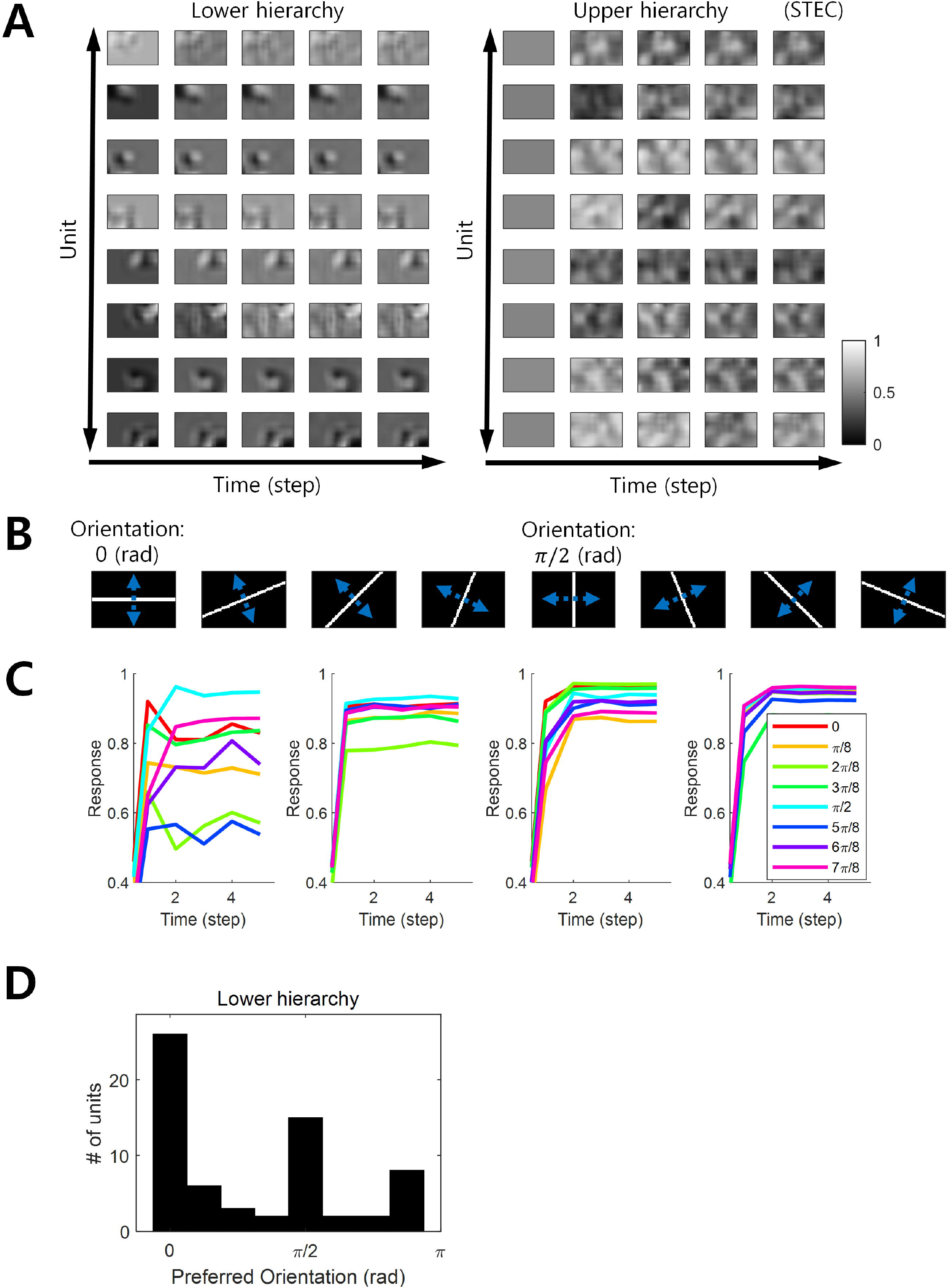
Orientation preference. (A) Examples of visual receptive fields of lower and upper hierarchy units. (B) Orientation images used to test the visual orientation preference of neuronal units at hierarchy 1 in the visual hierarchical structure that learned by the spatio-temporally efficient coding of natural scene images. White bars in each of the eight orientations are moved in the perpendicular directions denoted by blue dotted arrows. (C) Examples of lower hierarchy neural responses for bar stimuli. Each colour indicates each orientation. The neural responses of each orientation were obtained from the bar positions with the largest neural responses at time step 5. These four example units correspond to 1st, 3rd, 5th, and 7th units in (A). (D) Histograms of the orientation preference at time step.

## 4. Discussion

The present study aimed to find computational principles that enables visual hierarchical structures to attain the function to represent external visual information. To address the lack of neural coding principles to encompass both bottom-up and top-down pathways, we propose spatio-temporally efficient coding as a novel computational model. As a principled way of efficiently using given resources in both neural activity space and processing time, this coding principle optimises bidirectional information transmissions over hierarchical structures by simultaneously minimising temporally differences in neural responses and maximising entropy in neural representations. Simulation results showed that the proposed spatio-temporally efficient coding assigned the function of appropriate neural representations of natural scenes to a visual hierarchical structure on time domain and that it could predict deviations in neural responses to unlearned inputs and a bias in preferred orientations, which are well known characteristics of the visual system.

Appropriate neural representations in the present study were decodable stable neural representations which is an appropriate function to hierarchical brain structures on the time domain. To demonstrate this, we compared spatio-temporally efficient coding (STEC condition) with existing efficient coding (SEC condition) and sparse coding (Sparse condition) as shown in Fig 4. Only spatio-temporally efficient coding showed decodable stable neural representations. Existing predictive coding is conceptually problematic to apply to such real-time information processing (Hogendoorn and Burkitt, 2019). It would be interesting to compare the results with the conceptually improved predictive coding as well. Temporal coherence (Hurri and Hyvärinen, 2002; Zou et al., 2011) or slow feature analysis (Wiskott and Sejnowski, 2002; Berkes and Wiskott, 2005; Creutzig and Sprekeler, 2008) elicits smooth changes in neural responses, similar to temporally efficient coding in the present study. However, they were not used as comparative models because they are not intended to elicit different neural responses to different external inputs and are not suitable for direct application to the bidirectional multiple hierarchical structure of in the present study. Nevertheless, they can be substituted to the role of temporally efficient coding in the present study.

Since its initial proposal (Attneave, 1954; Barlow, 1961), spatially efficient coding has been validated experimentally (Laughlin, 1981). However, observed correlations between neurons, which maximise entropy to a lesser extent compared to mere spatially efficient coding assuming no inter-neuronal correlations, have yet to be incorporated into the principle of spatially efficient coding. Empirically observed neuronal correlations may drive computational processes of the brain away from strict spatially efficient coding. Recent studies suggest that biological visual systems are intermediate between strict spatially efficient coding and correlated neural responses (Stringer et al., 2019). Therefore, to create biologically plausible computational models, it is necessary to mitigate the spatially efficient coding objective by combining firing-rate-dependent correlations (de la Rocha et al., 2007). This enables more accurate information transmissions of visual perception mediated by visual hierarchical structures. As we focused on integrating spatially efficient coding with temporally efficient coding for computation in hierarchical structures, this study did not incorporate the correlations between neurons in spatially efficient coding, which will be pursued in follow-up studies.

Based on our simulations, we observed that the learning of bidirectional information transmission networks with spatio-temporally efficient coding was hindered when the balancing parameter *λ* was too small or too large (Fig 2C). Therefore, it was necessary to confine *λ* within a certain range, in which the magnitude of *λ* affected neural responses such that a larger *λ* rendered responses more variable (Fig 2C). Such increased variability is likely to originate from recurrent responses via higher hierarchies. This was confirmed by the observation that top-down synaptic weights become larger than bottom-up synaptic weights when *λ* increased (Fig 2B). Although a large *λ* attenuated the appropriateness of neural representations (Fig 3), it rendered stronger top-down synaptic connections in lower hierarchy (Fig 2B), which is consistent with the previous finding that top-down synaptic connections are stronger than bottom-up connections in the lateral geniculate nucleus (Sillito et al., 2006). As to why top-down synaptic weights increase with a larger *λ* value (Fig 2B), we speculate that learning via spatio-temporally efficient coding may increase the range of neural responses to maximise entropy through top-down pathways. While the bottom-up pathways originating from external inputs are invariant during learning, the top-down pathways originating from higher hierarchy neural responses are more flexible to adjustment to maximise entropy during learning. An increase in these top-down synaptic weights predicts impairment of eye movement tracking partially occluded visual targets in schizophrenic patients (Adams et al., 2012). This may be due to an increased higher hierarchy’s influence in patients with schizophrenia. The increased higher hierarchy’s influence in our simulations is that they do not stabilize neural responses sufficiently to distinguish a given input from other inputs (Fig 4C).

The deviant neural responses to unlearned inputs observed in this study (Fig 6) arise from smooth neural representations for learned inputs (distributed over the middle value). As such, neural representations for learned inputs extrude neural responses to unlearned inputs into a range of deviant neural responses. We conjecture that the visual system may generate deviant neural responses via a similar mechanism.

Spatio-temporally efficient coding predicted a bias in preferred orientations (Fig 8). In this regard, spatially efficient coding alone has been reported to predict bias in preferred orientations (Ganguli and Simoncelli, 2014). Notably, spatio-temporally efficient coding was able to predict this bias well, even when *λ* was low, that is, when the spatially efficient coding objective was less weighted (Fig 8D). Therefore, this bias prediction should be viewed as a result of spatio-temporally efficient coding, not as a result of spatially efficient coding alone.

The present study has several limitations. First, for simplicity, our simulation model contained only two hierarchies. However, it is necessary to explore how spatio-temporally efficient coding operates in models with more hierarchies. We also modelled 64 neuronal units at each hierarchy, as we assumed that this would be sufficient to represent the natural scene images used in this study. Nevertheless, the interactions between the number of neuronal units, levels of hierarchy, and spatio-temporally efficient coding require further investigation. Second, we demonstrated that the visual hierarchical structure could learn to represent static natural scene images with spatio-temporally efficient coding, but future follow-up studies will investigate whether the visual hierarchical structure learns to represent moving scenes using the same coding principle. Finally, the scope of the present study was limited to the visual system given that its hierarchical structure is well documented, but spatio-temporally efficient coding may be applied to other systems (e.g., somatosensory system) or to movements and planning.

### 4.1. Conclusions

In the present study, we proposed spatio-temporally efficient coding, inspired by the efficient use of given resources in neural systems, as a neural coding mechanism to assign representational functions to the hierarchical structures of the visual system. Simulations demonstrated that the visual hierarchical structure could represent the external world (i.e., natural scenes) appropriately using bidirectional information transmissions (Fig 2, 3, and 4). Furthermore, spatio-temporally efficient coding predicted the well-known properties of visual cortical neurons, including deviations in neural responses to unlearned images (Fig 6 and 7) and bias in preferred orientations (Fig 8). Our proposed spatio-temporally efficient coding may facilitate deeper mechanistic understanding of the computational processes of hierarchical brain structures.

## Data availability statement

All simulation and analysis codes are available at: https://github.com/DuhoSihn/Spatio-temporally-efficient-coding and on Zenodo (doi: 10.5281/zenodo.5298182)

## CRediT authorship contribution statement

**Duho Sihn**: Conceptualization, Methodology, Software, Validation, Formal analysis, Investigation, Resources, Data Curation, Writing - Original Draft, Writing - Review & Editing, Visualization, Supervision, Project administration. **Sung-Phil Kim**: Writing - Original Draft, Writing - Review & Editing, Visualization, Supervision, Project administration, Funding acquisition.

## Acknowledgments

This research was supported by the Brain Convergence Research Programs of the National Research Foundation (NRF) funded by the Korean government (MSIT) (NRF-2019M3E5D2A01058328 and No. 2021M3E5D2A01019542).

